# An untargeted metabolomics strategy to measure differences in metabolite uptake and excretion by mammalian cell lines

**DOI:** 10.1101/2020.06.02.129239

**Authors:** Marina Wright Muelas, Ivayla Roberts, Farah Mughal, Steve O’Hagan, Philip J. Day, Douglas B. Kell

**Author notes:** Correspondence to Marina Wright Muelas and Douglas B. Kell.

## Abstract

**Introduction:** It is widely but erroneously believed that drugs get into cells by passing through the phospholipid bilayer portion of the plasma and other membranes. Much evidence shows, however, that this is not the case, and that drugs cross biomembranes by hitchhiking on transporters for other natural molecules to which these drugs are structurally similar. Untargeted metabolomics can provide a method for determining the differential uptake of such metabolites.

**Objectives:** Blood serum contains many thousands of molecules and provides a convenient source of biologically relevant metabolites. Our objective was to measure them.

**Methods:** We develop an untargeted LC-MS/MS method to detect a broad range of compounds present in human serum. We apply this to the analysis of the time course of the uptake and secretion of metabolites in serum by several human cell lines, by analysing changes in the serum that represents the extracellular phase (the ‘exometabolome’ or metabolic footprint).

**Results:** Our method measures some 4,000-5,000 metabolic features in both ES^+^ and ES^−^ modes. We show that the metabolic footprints of different cell lines differ greatly from each other.

**Conclusion:** Our new, 15-minute untargeted metabolome method allows for the robust and convenient measurement of differences in the uptake of serum compounds by cell lines following incubation in serum, and its relation to differences in transporter expression.

## Introduction

One of the great unsolved problems of modern biology concerns the substrates of membrane transporters (Cesar-Razquin *et al.*, 2018; Cesar-Razquin *et al.*, 2015; Girardi *et al.*, 2020; Superti-Furga *et al.*, 2020), many of which remain ‘orphans’ (i.e. with unknown substrates), often despite the passage of decades since their identification in systematic genome-sequencing programmes (Ghatak *et al.*, 2019).

‘Untargeted metabolomics’ is a term nowadays commonly used to describe methods that seek the reproducible detection (and sometimes quantification) of small molecules in biological matrices (Cho *et al.*, 2014; Dunn *et al.*, 2013; Garg *et al.*, 2015; Martin *et al.*, 2015; Tautenhahn *et al.*, 2012; Treutler *et al.*, 2016). It is most commonly performed using chromatography coupled to mass spectrometry (e.g. (Dunn *et al.*, 2011; Dunn *et al.*, 2007; Dunn *et al.*, 2012)). A variety of methods, summarised in Table 1, have been developed for measuring the human serum metabolome, with both low- and high-resolution mass spectrometry. An ideal method would be rapid, reliable, and provide data on many (thousands of) metabolites simultaneously.

**Table 1.**
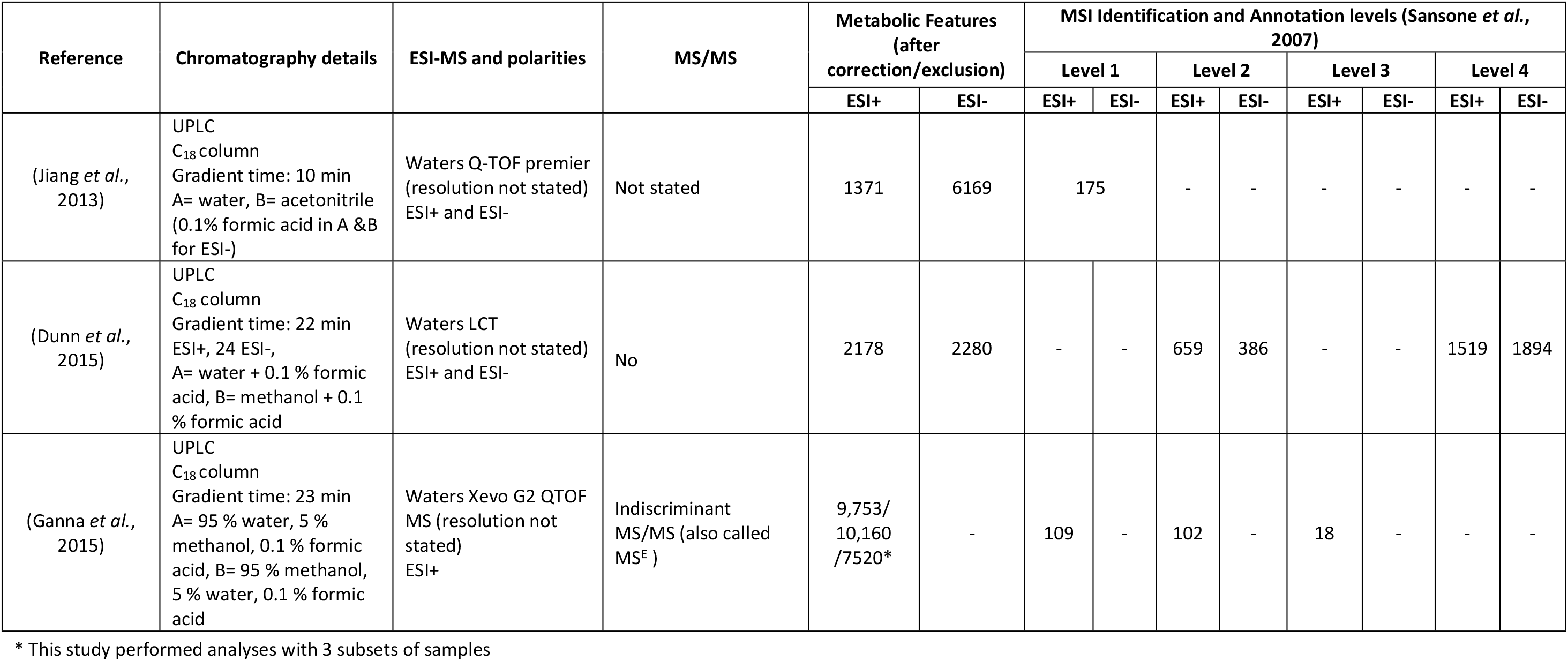
Selection of untargeted LC-MS studies of human serum. Note that only studies found where metabolic features as well as descriptions of metabolite identification levels have been included.

In a groundbreaking study using low-resolution LC-MS, Gründemann and colleagues (Gründemann *et al.*, 2005) recognised that incubation of cells containing different levels of a transporter of interest with plasma (as an reasonably unbiased source of candidate metabolites) might allow the discovery of transporter substrates by assessing their differential uptake into cells. They thereby discovered the transporter for the important nutraceutical (Borodina *et al.*, 2020) ergothioneine (Gründemann, 2012; Gründemann *et al.*, 2005).

As part of a wide-ranging study into the nature of transporter substrates (e.g. (Dobson and Kell, 2008; Jindal *et al.*, 2019; Kell *et al.*, 2013; Kell *et al.*, 2011; Kell and Oliver, 2014; Kell *et al.*, 2018)), we recognised that the approach of Gründemann and colleagues (Gründemann *et al.*, 2005) could be an ideal strategy for implementation using modern, high-resolution metabolomics methods. We have previously developed low-resolution methods for determining the human serum metabolome reliably (Broadhurst and Kell, 2006) and over extended periods (Begley *et al.*, 2009; Dunn *et al.*, 2011; Kenny *et al.*, 2010; Zelena *et al.*, 2009), including the extensive use of QA/QC samples. Our first requirement was thus to develop a new and robust method for untargeted serum metabolomics using modern, high-resolution instrumentation. The present paper describes this method, and an initial application to a series of human cell lines.

## Methods

### Cell culture

A549, K562, SAOS2 and U2OS cell lines were cultured in RPMI-1640 (Sigma, Cat No. R7509) culture media supplemented with 10 % fetal bovine serum (Sigma, Cat No. f4135) and 2 mM glutamine (Sigma, Cat No. G7513) without antibiotics. Cell cultures were maintained in T225 culture flasks (Star lab, CytoOne Cat No. CC7682-4225) kept in a 5% CO_2_ incubator at 37°C until 70-80 % confluent.

### Serum incubation experiments

#### Harvesting cells for serum incubation experiments

Cells from adherent cell lines were harvested by removing growth media and washing twice with 5 mL of pre-warmed Dulbecco’s Phosphate Buffered Saline (PBS) without calcium or magnesium (Gibco, Cat No. 14190094), then incubated in 3 mL of Gibco™ TrypLE™ Express Enzyme (1X), no phenol red (Gibco Cat No. 12604013) for 2-5 min at 37°C. At the end of incubation cells were resuspended in 5-7 mL of respective media when cells appeared detached to dilute TrypLE treatment. The cell suspension was transferred to 50 mL centrifuge tubes and immediately centrifuged at 300 x g for 5 min. Suspended cell lines were centrifuged directly from cultures in 50 mL centrifuge tubes and washed with PBS as above. The cell pellets were resuspended in 10-15 mL media and cell count and viability was determined using a Countess II FL Automated Cell Counter (ThermoFisher Scientific) set for Trypan Blue membrane exclusion method. Cells with > 95 % viability were used for serum incubation experiments.

#### Incubation of cells in serum

The procedure for incubation of cells in serum is described pictorially in Figure 1. More detailed information is provided in Supplementary Information file.

**Figure 1.**
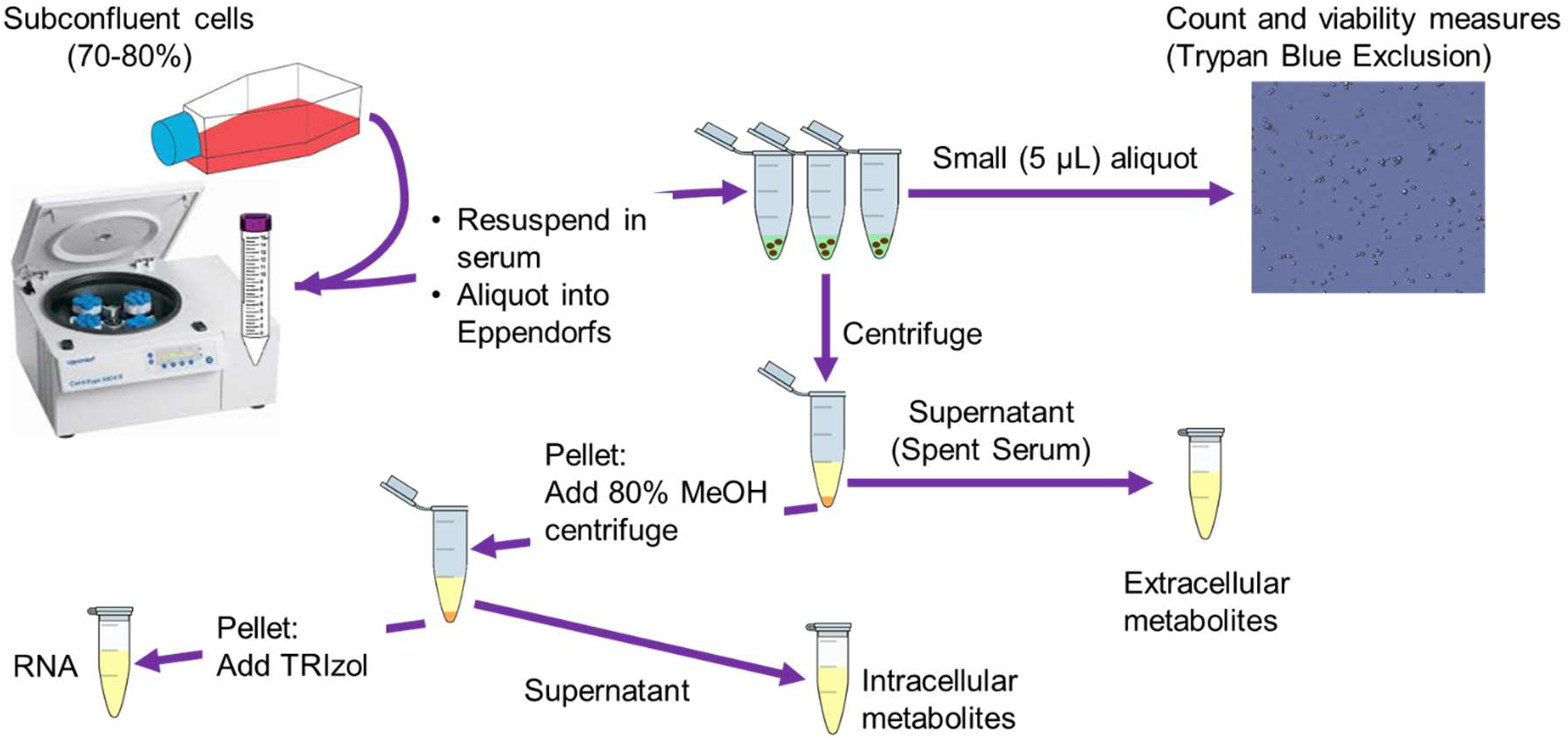
Incubation of cells in serum for metabolomics analysis to determine transporter substrates. Following incubation of cells in serum, spent serum is collected after centrifugation, followed by extraction using methanol. The remaining cell pellet is washed with PBS (at 37 °C), followed by quenching and extraction of intracellular metabolites using 80 % methanol. The spent medium and intracellular extracts are subsequently lyophilised (with a mixture of internal standards spiked in prior to lyophilisation) and reconstituted in water ready for analysis by LC-HRMS/MS.

### Internal standard solution mixture

An internal standard stock mixture was prepared using the following compounds and concentrations: citric acid-d4 (Cambridge Isotope Laboratories, DLM-3487), 225 μM; L-lysine-d_4_ (Cambridge Isotope Laboratories, DLM-2640), 112.5 μM; L-Tryptophan-(indole-d_5_) (Cambridge Isotope Laboratories, DLM-1092), 5.625 μM; stearic acid-d_35_ (Cambridge Isotope Laboratories, DLM-379), 225 μM; succinic acid-d_4_ (Sigma 293075), 112.5 μM; ^13^C_6_-carbamazepine (Sigma, C-136), 2.25 μM; Leucine-d_10_ (Sigma 492949), 22.5 μM and methionine-d_4_ (Cambridge Isotope Laboratories, DLM-2933), 22.5 μM.

### Sample preparation for metabolomics analysis

Fresh or spent serum samples were thawed at room temperature and maintained on ice throughout the sample preparation process. Samples were prepared by addition of 100 μL sample to a 2 mL Eppendorf containing 330 μL Methanol (LC-MS grade) and 20 μL of internal standards mix (ISTDs). The mixture of methanol and ISTDs was previously cooled at −80 °C and maintained on dry ice when adding serum. The mixture of serum, methanol and ISTD was vortexed vigorously followed by centrifugation at 13,300 rpm for 15 min at 4 °C to pellet proteins. Multiple 75 μL aliquots (for extraction replicates) of the resulting supernatant dried in a vacuum centrifuge (ScanVac MaxiVac Beta Vacuum Concentrator system, LaboGene ApS, Denmark) with no temperature application and stored at −80 °C until required for LC-MS/MS analysis.

Quality controls (QC) and conditioning QC samples were also prepared in this way using pooled human serum.

Extraction blanks were prepared in the same way as spent serum samples replacing serum and internal standard mix with 120 μL of water (LC-MS grade). Evaluation/system suitability samples were also prepared by replacing serum with water.

Prior to analysis, samples were resuspended in 40 μL water (LC-MS), centrifuged at 13,300 rpm for 15 min at 4 °C to remove any particulates and transferred to glass sample vials.

### HPLC-MS /MS analysis of spent serum samples

Untargeted HPLC-MS/MS data acquisition was performed following methodologies and guidelines in (Broadhurst *et al.*, 2018; Broadhurst and Kell, 2006; Brown *et al.*, 2005; Dunn *et al.*, 2011; Mullard *et al.*, 2015). Data were acquired using a ThermoFisher Scientific Vanquish HPLC system coupled to a ThermoFisher Scientific Q-Exactive mass spectrometer (ThermoFisher Scientific, UK). A resolution of 70,000 was used for MS and 17,500 for ddMS (further details provided in Supplementary Information). Chromatographic separation was performed on a Hypersil Gold aQ column (C18 2.1 mm x 100 mm, 1.9 μm, ThermoFisher Scientific) operating at a column temperature of 50°C. Elution was performed over 15 minutes at a flow rate of 0.4 mL/min using two solvents: 0.1 % formic acid in water (solvent A) and 0.1 % formic acid in methanol (solvent B). as described in Table 2 below. Column eluent was diverted to waste in the first 0.4 minutes and the last 0.1 minutes of the gradient. Sample vials were stored at 4°C in the HPLC autosampler, with 5 μL injected for positive ionisation and 15 μL for negative ionisation. MS acquisition settings are described in supplementary information 1.

**Table 2.** HPLC gradient elution program applied for HPLC-MS/MS analysis for ESI+ and ESI− mode

| Time (minutes) | Flow rate (mL/min) | Solvent A (%) | Solvent B (%) |
| --- | --- | --- | --- |
| 0 | 0.4 | 99.0 | 1.0 |
| 2 | 0.4 | 99.0 | 1.0 |
| 10 | 0.4 | 1.0 | 99.0 |
| 12 | 0.4 | 1.0 | 99.0 |
| 13 | 0.4 | 99.0 | 1.0 |
| 15 | 0.4 | 99.0 | 1.0 |

Samples were analysed following guidelines set out in (Dunn *et al.*, 2011) and (Broadhurst *et al.*, 2018). Briefly, blank extraction samples were injected at the beginning and end of each batch to assess carry over and lack of contamination. Isotopically labelled internal standards were added to analytical and QC samples to assess system stability throughout the batch. QC samples, prepared with a standard reference material, pooled human serum, were also applied to condition the analytical platform, enable reproducibility measurements and to correct for systematic errors.

### LC-MS/MS data preprocessing and analysis

Raw instrument data (.RAW) were exported to Compound Discoverer 3.1 (CD3.1) for deconvolution, alignment, QC based area correction and annotation (full workflow and settings are provided in supplementary information 2).

For analysis of data for serum compound uptake and excretion by cell lines, peak areas from CD3.1 were exported as an excel file into a KNIME workflow developed in-house for data analysis and visualisation (available on request). Within this workflow, Principal Components Analysis was used to visualise trends. Subsequently, simple univariate statistical analyses were carried out on log_2_ transformed data using a paired t-test. Volcano plots were created using these data, with a threshold of P < 0.05 and absolute log_2_ fold change > 0.5 set for defining a notable change in compound abundance between time points compared. UpSet plots (Lex *et al.*, 2014) (Supplementary Figure 3) were then used to find unique and shared consumed or secreted compounds between cell lines.

For all data acquired, annotation and identification criteria were according to (Schymanski *et al.*, 2014).

## Results

### A RP-LC-ESI-MS/MS method capable of detecting a broad range of serum compounds

The optimisation of chromatographic (O’Hagan *et al.*, 2005) and mass spectrometric (Vaidyanathan *et al.*, 2003) methods typically requires trade-offs between multiple objectives. During the development of the LC-MS/MS method used here our aim was both to maximise the number of serum compounds detected but also to acquire sufficient fragmentation data for more confident identification, with both achieved within a reasonably short period. Table 3 shows a summary of the results obtained from CD3.1 using the data acquired using the LC-MS/MS method described in both ESI+ and ESI− modes after running a set of serum QC samples. The method we have developed enables the detection of a large number of metabolic features, of which around 65 % can be attributed to sample-related compounds (after background subtraction and exclusion, removal of compounds not found in > 50 % of QC samples, and exclusion of compounds with a QC CV > 30%, see supplementary information 2). Around 80 % or greater of these sample compounds had a QC CV < 15 % demonstrating a good level of reproducibility across injections. As can be seen in Supplementary Figure 2 peak areas of detected spiked internal standards displayed high reproducibility (<10% RSD) and excellent mass accuracy (< 1ppm) across QC injections throughout the run.

**Table 3.**
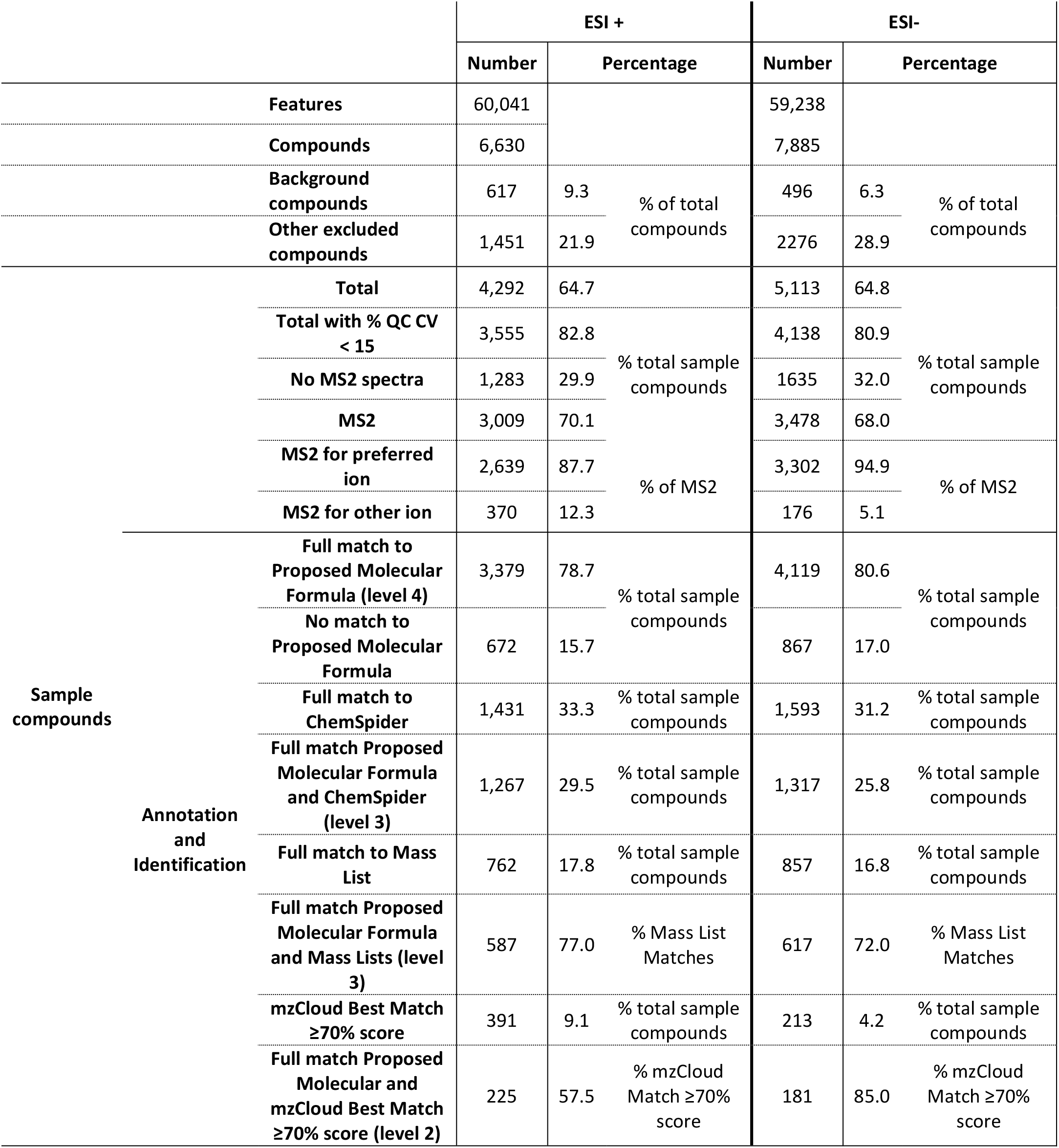
Summary of LC-MS/MS results of serum QC samples obtained following preprocessing using CD3.1. Note features here correspond to detected ions, compounds here correspond to what is commonly referred to as metabolic features (m/z and RT).

To improve confidence in metabolite annotation we also performed data dependent MS/MS across a range of masses in a similar way to (Mullard *et al.*, 2015). As can be seen in Table 3 around 70 % of sample compounds had associated MS/MS spectra with a large proportion of MS2 spectra corresponding to a preferred adduct ion ([M+H]+).

As is commonly the case in untargeted metabolomics, the level of annotation and identification confidence was quite varied. In our analyses, annotation and identification ranged from identification levels 5-2 (Schymanski *et al.*, 2014) (Table 3). We do find a small number of level 1 identifications in our study; however, our in-house mass and spectral libraries are small and still under development. For this reason, level 1 identifications will not be discussed. Results in terms of annotation and identification at various levels from both ESI+ and ESI-were comparable (Table 2); we will describe these for ESI+.

In ESI+ 3,376 sample compounds had a full match to a proposed molecular formula (level 4). Of these, 1,010 matched mass libraries in ChemSpider (level 3) on our searches against BioCyc, HMDB, KEGG, MassBank and NIST. In addition, nearly 700 compounds were matched against mass libraries provided by CD3.1 software along with our own imported ones (including the serum metabolome database (Psychogios *et al.*, 2011) and the COCONUT Natural Products database (Sorokina and Steinbeck, 2020)). Of the 3,009 compounds with MS/MS spectra, 391 were matched with reasonable confidence (≥ 70%) to the mzCloud spectral library. Of these, 226 were fully matched against a proposed molecular formula by CD3.1 providing level 2 identification confidence. These compounds represented diverse metabolite classes such as amino acids, peptides and analogues, lipids and lipid-like molecules (including fatty acyls and steroids), as well as carboxylic acids and derivatives.

Of the remaining 165 compounds with ≥ 70% match to the mzCloud spectral library but without full match to a proposed molecular formula, a large proportion are not marked as having a full match to proposed molecular formulae for several reasons. This may include better spectral matching with other compounds sharing similar substructures, but with this providing a mass error > 3 ppm. In some cases, closer manual inspection is required whereas in others, the spectral matching provides clues as to the possible underlying substructure of the molecule in question. There is some level of duplication in annotation and putative identifications, either through close matches to mass or spectral libraries, or due to the same precursor mass appearing at more than one retention time (yet with different peak areas and intensities). As an example, we found Arachidonic acid eluting at 4 different retention times yet spectral matching of all these with mzCloud was > 90 % in confidence. Another such example was Ecgonine, eluting at two different retention times yet both having excellent spectral matching to mzCloud (> 90 % match). Further investigation may reveal this to be due to (positional) isomers and/or some compounds simply binding to the column and eluting over different times. This is of lesser importance for present purposes where we aim to find unique markers of transporter substrates.

Taking into account that these results are illustrative of a QC run, results in Table 3 suggests our LC-MS/MS method is a clear improvement on those shown in Table 1. The number of metabolic features after correction and exclusion in our study are lower than those in Ganna et al (Ganna *et al.*, 2015), however, the number of level 2 identified compounds (whether using (Sumner *et al.*, 2007) or (Schymanski *et al.*, 2014)) is nearly double. Whilst this is not the case when we compare against the number of level 2 identified compounds in (Dunn *et al.*, 2015), we expect this will increase in our methodology as number of compounds and spectra in mzCloud grow, we match to spectral libraries outside of Compound Discoverer and we continue to increase our in-house library. Furthermore, our elution gradient is shorter (15 min in both ESI+ and ESI-vs 22 min ES+ and 24 min ESI−) which will provide great time and cost savings.

### Application of our method to determine the consumption and excretion of serum compounds by mammalian cell lines

One of the main purposes for developing the above untargeted LCMSMS methodology is to apply this to measure the uptake and excretion of serum compounds by mammalian cell lines. As a proof of principle, we assessed this by analysis in ESI+ of spent serum samples from 4 cell lines (A549, K562, SAOS2 and U2OS) incubated in serum at two different densities (2 and 4 million) at two timepoints (0 and 20 minutes incubation).

A summary table of the number of compounds detected and identified with various levels of confidence can be found in Supplementary Table S1. The results obtained are comparable to those shown in Table 1, demonstrating good reproducibility of our method.

PCA reveals differences in samples (Figure 2): the different cell lines fell into distinct groups separated in both PC1 and PC2. Furthermore, separations between time 0 and 20 mins of incubation suggested differences in metabolic profile of spent serum influenced by time.

**Figure 2.**
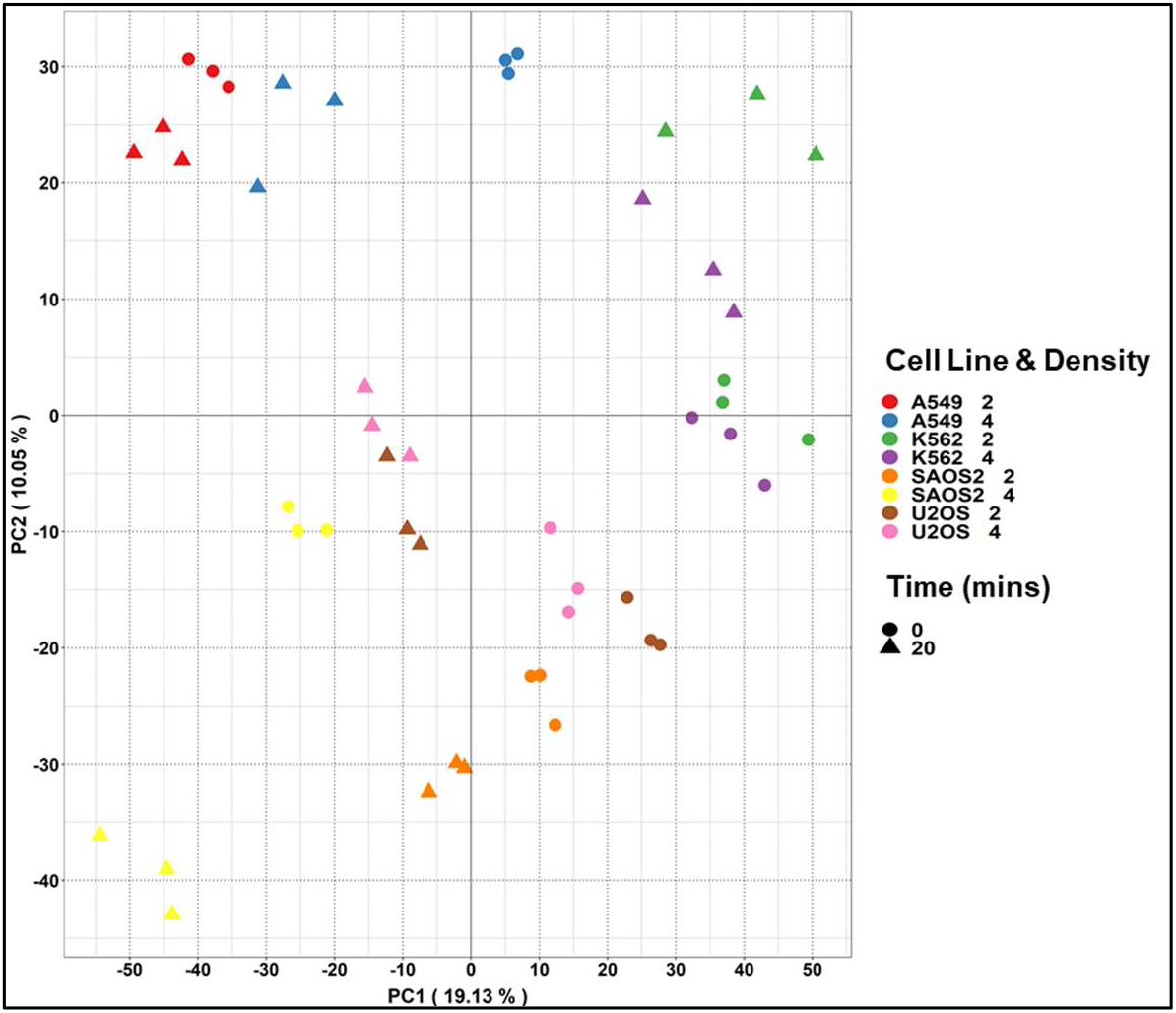
PCA scores plots of spent serum extracts following incubation for 0 or 20 minutes with 4 different cell lines and two densities. Colour: Cell line and density, shape: incubation time.

To confirm that the separation of cell lines as well as the effect of incubation time and cell density are the result of differences in the uptake and excretion of serum components, we performed simple univariate analyses. The volcano plots in Figure 3 confirm this is the case; cell lines consumed and excreted serum components however, the number of compounds consumed or excreted was not always proportional to increasing density, indicating that a rich set of metabolic activities were taking place during this period.

**Figure 3.**
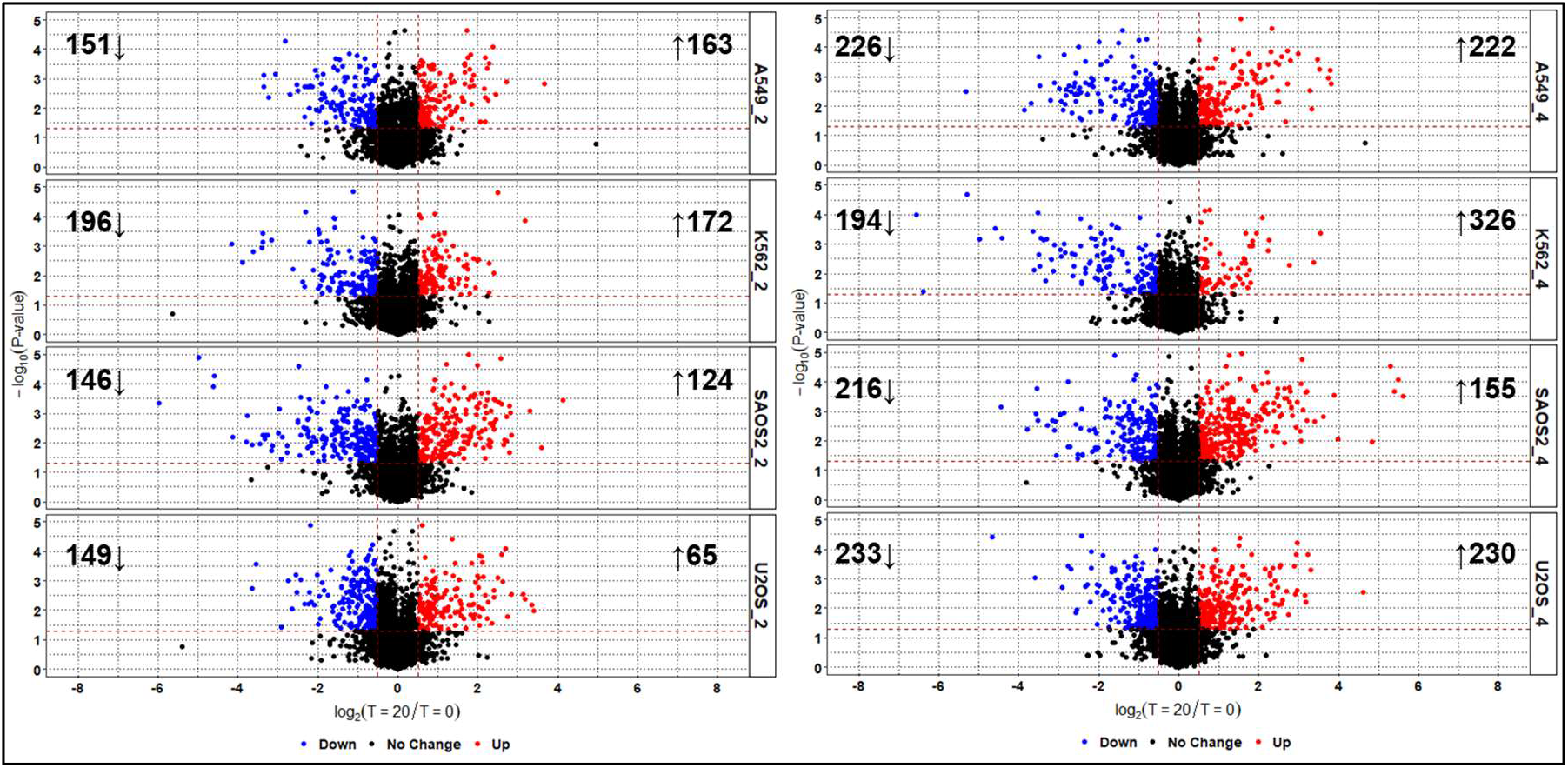
Volcano plots showing differences in number and magnitude of serum compound consumption and excretion by different cell lines and densities over 2 minutes. Threshold for significant change: P-value < 0.05 and log_2_ Fold Change < −0.5 or > 0.5. Left panel: 2 million cell density; right side 4 million.

Fuller results will be reported elsewhere for a much larger panel of cell lines; however, we provide examples of a compound exclusively consumed by one cell line (SAOS2, Figure 4A), another exclusively secreted by another cell line (A549, Figure 4B) and finally one where a mix of consumption and secretion was observed (Figure 4C).

**Figure 4.**
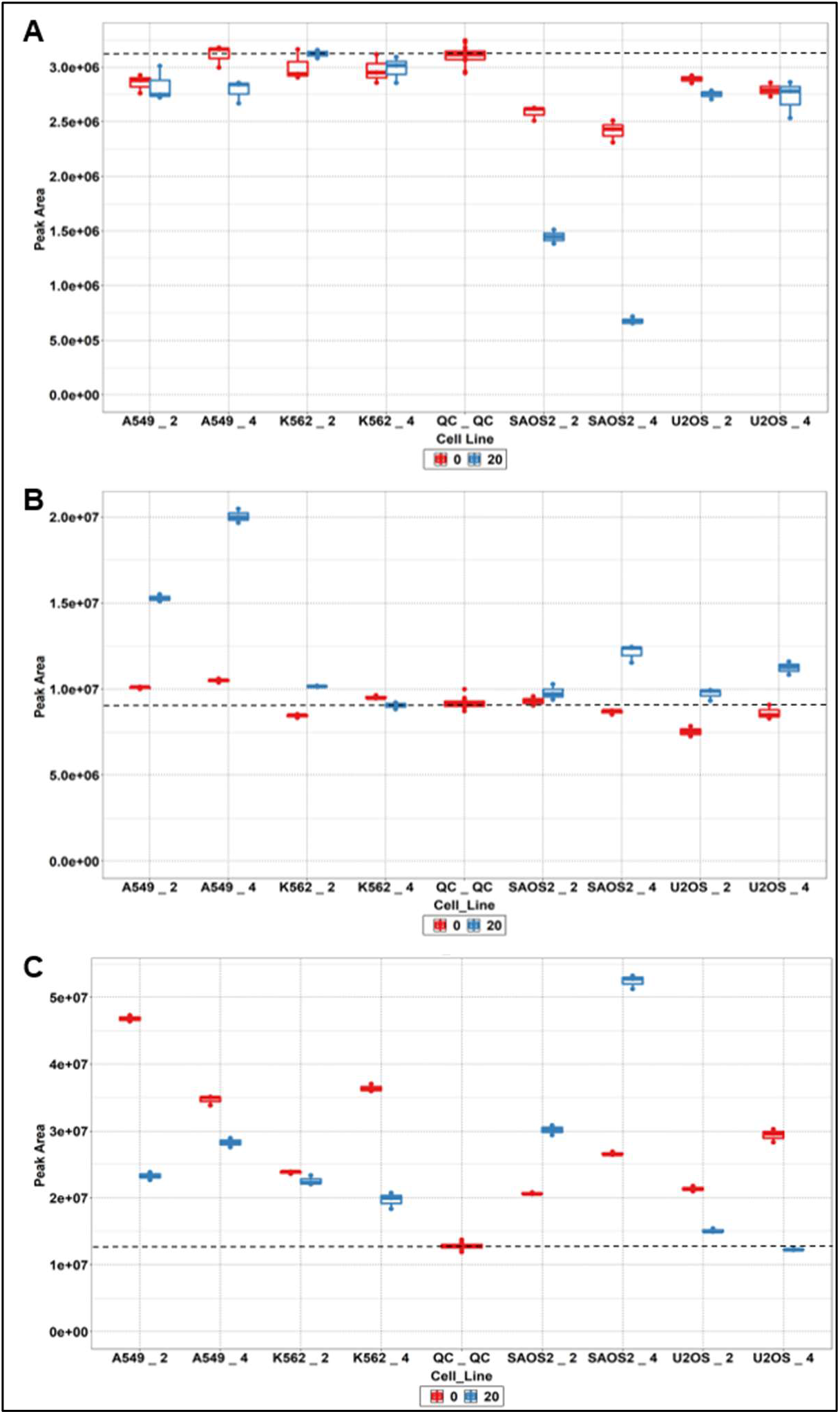
Cell-line specific consumed or secreted compounds A) Unidentified compound with assigned molecular formula C21H37N5O7 consumed by SAOS2 cell lines only, B) γ-L-Glutamyl-L-glutamic acid secreted exclusively by A549 cell lines. C) Nicotinamide secreted by SAOS2 cell lines but consumed by others. X-axis labels: A549_2, A549 cells at 2 million density; A549_4, A549 cells at 4 million density; K562_2, K562 cells at 2 million density; A549_4, K562 cells at 4 million density; QC_QC, Quality Control; SAOS2_2, SAOS2 cells at 2 million density; SAOS2_4, SAOS2 cells at 4 million density; U2OS _2, U2OS cells at 2 million density; U2OS_4, U2OS cells at 4 million density. Horizontal dashed line added at the median level in QC sample to aid visualisation.

In SAOS2 cells, an unknown compound with mass of 471.26882 matching the molecular formula C21H37N507 was consumed exclusively by this cell lines, with consumption increasing in relation to higher cell density (Figure 4A, log_2_ fold change −0.84, P = 0.0004 at 2 million density and log2 fold change −1.83, P = 0.0002 at 4 million density). Fragmentation data matching against mzCloud suggest some possible substructures of this compound match fragments for an environmental compound on the NORMAN suspect list (Mistrik *et al.*, 2019) as shown in Supplementary Figure 3A.

As can be seen in Figure 4B, A549 cells secreted γ-L-Glutamyl-L-glutamic acid whilst the other 3 cell lines did not. The increase in levels of this metabolite was nearly double when cell density was doubled (log_2_ fold change 0.60, P = 0.0004 at 2 million density and log_2_ fold change 0.93, P = 0.0002 at 4 million density) whereas in other cell lines the changes were below our threshold (log_2_ fold change > 0.5 and P < 0.05). The identification of this compound is at a reasonable level 2, with 90.2 % match to this compound in mzCloud, and low mass error (−0.00029 Da or −1.06 ppm) as can be seen in Supplementary Figure 3B.

Nicotinamide (level 2 identification as shown in Supplementary Figure 3C) was found to be secreted by SAOS2 cell lines in a density dependent manner (log_2_ fold change 0.55, P = 0.0010 at 2 million density and log2 fold change 0.98, P = 0.002 at 4 million density) yet consumed by K562 (log_2_ fold change −0.89, P = 0.0050) and U2OS (log_2_ fold change −1.26, P = 0.0004) cell lines at 4 million density and A549 cells at 2 million density only (log_2_ fold change of −1.01, P = 0.0002).

While the biological significance of the consumed or secreted compounds in Figure 3 and Figure 4 will be discussed elsewhere in due course, the main message from this study is that the methodology employed here is robust and reproducible, (ii) capable of measuring the transport behaviour of serum compounds in an entirely unbiased way, (iii) and shows the massive differences between individual cell lines (O’Hagan *et al.*, 2018; Wright Muelas *et al.*, 2019).

## Discussion

We have here described an untargeted LC-MS/MS method developed to maximise the number and diversity of compounds detected in human serum whilst also acquiring sufficient fragmentation data for improved metabolite annotation confidence, all within a reasonable period of 15 minutes. The method enables detection of around 4,000-5,000 sample-related metabolic features in both ESI+ and ESI−, with excellent reproducibility and mass accuracy; across QC injections, ≥ 80 % of sample compounds QC CVs were ≤ 15 % (Table 3), and spiked internal standard QC CVs were < 10 % (Supplementary Figure S2) with excellent mass accuracy (< 1ppm).

Annotation and identification of metabolites is by far the greatest bottleneck encountered in untargeted metabolomics (Djoumbou Feunang *et al.*, 2016; Dunn *et al.*, 2013; Misra and van der Hooft, 2016). Despite an increasing availability of mass spectral libraries (Vinaixa *et al.*, 2016), only a small proportion of small molecules in these are derived from experimental data using pure standards and, even then, these seem to cover only around 40% of compounds within human genome scale metabolic network reconstructions (Frainay *et al.*, 2018). Note that most serum metabolites have an exogenus source(O’Hagan and Kell, 2017). Only a limited number of untargeted metabolomics studies of human serum using LC-MS/MS have sufficient details with which to compare our results (Table 1). Our method enables annotation of a significant number of metabolites at levels 4-2 (Table 3). From these, we found 226 metabolites with level 2 identification confidence in ESI+, representing diverse (and relevant) metabolite classes. The number of level 2 identified compounds using our method is also improved in comparison to results reported by Ganna et al (Ganna *et al.*, 2015). This is not the case when we compared (Dunn *et al.*, 2015), however, the elution gradient in our method is shorter (15 min in both ESI+ and ESI− vs 22 min ES+ and 24 min ESI−) which will provide great time and cost savings. The annotation and identification of metabolites to level 2 in our study is likely also limited by the use of spectral libraries available through Compound Discoverer, namely mzCloud and local spectral libraries provided as standard with this software within mzVault. Another limitation within our reported data is the small number of level 1 metabolite identifications in our study; our in-house mass and spectral libraries are small and still under development.

In addition to maximising metabolite detection and identification, we have demonstrated the applicability of the LC-MS/MS method described to measure differences in the uptake and secretion of compounds by cell lines following incubation in human serum. This takes inspiration from work by Gründemann and colleagues (Gründemann *et al.*, 2005) taking advantage of the complex mixture of candidate transporter substrates in human serum. The results reveal both the reproducibility of the analyses and distinct metabolic footprints(Allen *et al.*, 2003) of different cell lines in terms of both uptake and secretion (Figures 2, 3).

As shown in Figure 4, some compounds were consumed exclusively by certain cell lines and not others, whilst others were consumed by some but secreted by others. These differences are undoubtedly related to the transporter expression profiles of these cell lines. We have recently shown transporter expression to vary widely between cells and tissues which can be explained by the requirements of different tissues and cell lines for different amounts of specific substrates (O’Hagan *et al.*, 2018). Fuller and more extensive results using a larger panel of different cell lines and time points will be reported elsewhere, and use of transcriptomic and proteomic transporter expression profiles to relate these to the potential substrates of transporter proteins.

## Conclusion

We have developed a new, 15-minute untargeted metabolomics method using LC-MS/MS that allows for the robust and convenient measurement of a large number of metabolites in human serum. The method additionally acquires fragmentation data to enable improved annotation and identification of compounds. We also describe a protocol for investigating the natural substrates of transporters by way of incubating human cell lines in serum and using the above LC-MS/MS method to measure reproducible and unbiased differences in the uptake of serum compounds.

## Supporting information

Supplementary Information 2: Compound Discoverer settings

## Acknowledgements

We thank the BBSRC (grant BB/P009042/1) and the Novo Nordisk Foundation (grant NNF10CC1016517) for financial support.

## Author Information

### Contributions

D.B.K. and P.J. D. highlighted the utility of the use of serum as a medium to incubate cells in to measure differences in uptake and excretion as described in (Gründemann *et al.*, 2005), and obtained funding for the study. M.W.M. designed cell culture experiments and developed LC-MS/MS analyses described. F.M., M.W.M. and I.G. performed cell culture and uptake experiments. M.W.M. and I.G. performed sample preparation and LC-MS/MS analysis. M.W.M. analysed data. SOH contributed to KNIME workflows for data analysis. D.B.K and M.W.M. wrote the manuscript. All authors contributed to the writing and approval of the manuscript.

## Ethics declarations

### Ethical approval

This article does not contain any studies with human and/or animal participants performed by any of the authors.

**Data Availability Statement**

## Electronic Supplementary Material

Supplementary Information 1: Supplementary Methods, Figures and Tables

Supplementary Information 2: Compound Discoverer settings

## References

Allen, J., Davey, H.M., Broadhurst, D., Heald, J.K., Rowland, J.J., Oliver, S.G. and Kell, D.B. (2003) High-throughput classification of yeast mutants for functional genomics using metabolic footprinting. Nature Biotechnology 21, 692–696.

Begley, P., Francis-McIntyre, S., Dunn, W.B., Broadhurst, D.I., Halsall, A., Tseng, A., Knowles, J., Goodacre, R. and Kell, D.B. (2009) Development and Performance of a Gas Chromatography−Time-of-Flight Mass Spectrometry Analysis for Large-Scale Nontargeted Metabolomic Studies of Human Serum. Analytical Chemistry 81, 7038–7046.

Borodina, I., Kenny, L.C., McCarthy, C.M., Paramasivan, K., Pretorius, E., Roberts, T.J., van der Hoek, S.A. and Kell, D.B. (2020) The biology of ergothioneine, an antioxidant nutraceutical. Nutrition Research Reviews, 1–28.

Broadhurst, D., Goodacre, R., Reinke, S.N., Kuligowski, J., Wilson, I.D., Lewis, M.R. and Dunn, W.B. (2018) Guidelines and considerations for the use of system suitability and quality control samples in mass spectrometry assays applied in untargeted clinical metabolomic studies. Metabolomics 14, 72.

Broadhurst, D.I. and Kell, D.B. (2006) Statistical strategies for avoiding false discoveries in metabolomics and related experiments. Metabolomics 2, 171–196.

Brown, M., Dunn, W.B., Ellis, D.I., Goodacre, R., Handl, J., Knowles, J.D., O’Hagan, S., Spasić, I. and Kell, D.B. (2005) A metabolome pipeline: from concept to data to knowledge. Metabolomics 1, 39–51.

Cesar-Razquin, A., Girardi, E., Yang, M., Brehme, M., Saez-Rodriguez, J. and Superti-Furga, G. (2018) In silico Prioritization of Transporter-Drug Relationships From Drug Sensitivity Screens. Front Pharmacol 9, 1011.

Cesar-Razquin, A., Snijder, B., Frappier-Brinton, T., Isserlin, R., Gyimesi, G., Bai, X., Reithmeier, R.A., Hepworth, D., Hediger, M.A., Edwards, A.M. and Superti-Furga, G. (2015) A Call for Systematic Research on Solute Carriers. Cell 162, 478–87.

Cho, K., Mahieu, N.G., Johnson, S.L. and Patti, G.J. (2014) After the feature presentation: technologies bridging untargeted metabolomics and biology. Current Opinion in Biotechnology 28, 143–148.

Djoumbou Feunang, Y., Eisner, R., Knox, C., Chepelev, L., Hastings, J., Owen, G., Fahy, E., Steinbeck, C., Subramanian, S., Bolton, E., Greiner, R. and Wishart, D.S. (2016) ClassyFire: automated chemical classification with a comprehensive, computable taxonomy. Journal of Cheminformatics 8, 61.

Dobson, P.D. and Kell, D.B. (2008) Carrier-mediated cellular uptake of pharmaceutical drugs: an exception or the rule? Nat Rev Drug Discov 7, 205–20.

Dunn, W.B., Broadhurst, D., Begley, P., Zelena, E., Francis-McIntyre, S., Anderson, N., Brown, M., Knowles, J.D., Halsall, A., Haselden, J.N., Nicholls, A.W., Wilson, I.D., Kell, D.B., Goodacre, R. and Human Serum Metabolome, C. (2011) Procedures for large-scale metabolic profiling of serum and plasma using gas chromatography and liquid chromatography coupled to mass spectrometry. Nat Protoc 6, 1060–83.

Dunn, W.B., Broadhurst, D.I., Deepak, S.M., Buch, M.H., McDowell, G., Spasic, I., Ellis, D.I., Brooks, N., Kell, D.B. and Neyses, L. (2007) Serum metabolomics reveals many novel metabolic markers of heart failure, including pseudouridine and 2-oxoglutarate. Metabolomics 3, 413–426.

Dunn, W.B., Erban, A., Weber, R.J.M., Creek, D.J., Brown, M., Breitling, R., Hankemeier, T., Goodacre, R., Neumann, S., Kopka, J. and Viant, M.R. (2013) Mass appeal: metabolite identification in mass spectrometry-focused untargeted metabolomics. Metabolomics 9, 44–66.

Dunn, W.B., Lin, W., Broadhurst, D., Begley, P., Brown, M., Zelena, E., Vaughan, A.A., Halsall, A., Harding, N., Knowles, J.D., Francis-McIntyre, S., Tseng, A., Ellis, D.I., O’Hagan, S., Aarons, G., Benjamin, B., Chew-Graham, S., Moseley, C., Potter, P., Winder, C.L., Potts, C., Thornton, P., McWhirter, C., Zubair, M., Pan, M., Burns, A., Cruickshank, J.K., Jayson, G.C., Purandare, N., Wu, F.C.W., Finn, J.D., Haselden, J.N., Nicholls, A.W., Wilson, I.D., Goodacre, R. and Kell, D.B. (2015) Molecular phenotyping of a UK population: defining the human serum metabolome. Metabolomics 11, 9–26.

Dunn, W.B., Wilson, I.D., Nicholls, A.W. and Broadhurst, D. (2012) The importance of experimental design and QC samples in large-scale and MS-driven untargeted metabolomic studies of humans. Bioanalysis 4, 2249–64.

Frainay, C., Schymanski, E.L., Neumann, S., Merlet, B., Salek, R.M., Jourdan, F. and Yanes, O. (2018) Mind the Gap: Mapping Mass Spectral Databases in Genome-Scale Metabolic Networks Reveals Poorly Covered Areas. Metabolites 8, 51.

Ganna, A., Fall, T., Salihovic, S., Lee, W., Broeckling, C.D., Kumar, J., Hägg, S., Stenemo, M., Magnusson, P.K.E., Prenni, J.E., Lind, L., Pawitan, Y. and Ingelsson, E. (2015) Large-scale non-targeted metabolomic profiling in three human population-based studies. Metabolomics 12, 4.

Garg, N., Kapono, C.A., Lim, Y.W., Koyama, N., Vermeij, M.J.A., Conrad, D., Rohwer, F. and Dorrestein, P.C. (2015) Mass spectral similarity for untargeted metabolomics data analysis of complex mixtures. International Journal of Mass Spectrometry 377, 719–727.

Ghatak, S., King, Z.A., Sastry, A. and Palsson, B.O. (2019) The y-ome defines the 35% of Escherichia coli genes that lack experimental evidence of function. Nucleic Acids Research 47, 2446–2454.

Girardi, E., César-Razquin, A., Lindinger, S., Papakostas, K., Konecka, J., Hemmerich, J., Kickinger, S., Kartnig, F., Gürtl, B., Klavins, K., Sedlyarov, V., Ingles-Prieto, A., Fiume, G., Koren, A., Lardeau, C.-H., Kumaran Kandasamy, R., Kubicek, S., Ecker, G.F. and Superti-Furga, G. (2020) A widespread role for SLC transmembrane transporters in resistance to cytotoxic drugs. Nature Chemical Biology 16, 469–478.

Gründemann, D. (2012) The ergothioneine transporter controls and indicates ergothioneine activity — A review. Preventive Medicine 54, S71–S74.

Gründemann, D., Harlfinger, S., Golz, S., Geerts, A., Lazar, A., Berkels, R., Jung, N., Rubbert, A. and Schömig, E. (2005) Discovery of the ergothioneine transporter. Proc Natl Acad Sci U S A 102, 5256–61.

Jiang, M., Chen, T., Feng, H., Zhang, Y., Li, L., Zhao, A., Niu, X., Liang, F., Wang, M., Zhan, J., Lu, C., He, X., Xiao, L., Jia, W. and Lu, A. (2013) Serum Metabolic Signatures of Four Types of Human Arthritis. Journal of Proteome Research 12, 3769–3779.

Jindal, S., Yang, L., Day, P.J. and Kell, D.B. (2019) Involvement of multiple influx and efflux transporters in the accumulation of cationic fluorescent dyes by *Escherichia coli*. bioRxiv, 603688.

Kell, D.B., Dobson, P.D., Bilsland, E. and Oliver, S.G. (2013) The promiscuous binding of pharmaceutical drugs and their transporter-mediated uptake into cells: what we (need to) know and how we can do so. Drug Discov Today 18, 218–39.

Kell, D.B., Dobson, P.D. and Oliver, S.G. (2011) Pharmaceutical drug transport: the issues and the implications that it is essentially carrier-mediated only. Drug Discovery Today 16, 704–714.

Kell, D.B. and Oliver, S.G. (2014) How drugs get into cells: tested and testable predictions to help discriminate between transporter-mediated uptake and lipoidal bilayer diffusion. Frontiers in Pharmacology 5.

Kell, D.B., Wright Muelas, M., O’Hagan, S. and Day, P.J. (2018) The role of drug transporters in phenotypic screening. Drug Target Review, 16–19.

Kenny, L.C., Broadhurst, D.I., Dunn, W., Brown, M., North, R.A., McCowan, L., Roberts, C., Cooper, G.J.S., Kell, D.B. and Baker, P.N. (2010) Robust Early Pregnancy Prediction of Later Preeclampsia Using Metabolomic Biomarkers. Hypertension 56, 741–749.

Lex, A., Gehlenborg, N., Strobelt, H., Vuillemot, R. and Pfister, H. (2014) UpSet: Visualization of Intersecting Sets. IEEE Transactions on Visualization and Computer Graphics 20, 1983–1992.

Martin, J.-C., Maillot, M., Mazerolles, G., Verdu, A., Lyan, B., Migné, C., Defoort, C., Canlet, C., Junot, C., Guillou, C., Manach, C., Jabob, D., Bouveresse, D.J.-R., Paris, E., Pujos-Guillot, E., Jourdan, F., Giacomoni, F., Courant, F., Favé, G., Le Gall, G., Chassaigne, H., Tabet, J.-C., Martin, J.-F., Antignac, J.-P., Shintu, L., Defernez, M., Philo, M., Alexandre-Gouaubau, M.-C., Amiot-Carlin, M.-J., Bossis, M., Triba, M.N., Stojilkovic, N., Banzet, N., Molinié, R., Bott, R., Goulitquer, S., Caldarelli, S. and Rutledge, D.N. (2015) Can we trust untargeted metabolomics? Results of the metabo-ring initiative, a large-scale, multi-instrument inter-laboratory study. Metabolomics 11, 807–821.

Misra, B.B. and van der Hooft, J.J.J. (2016) Updates in metabolomics tools and resources: 2014–2015. ELECTROPHORESIS 37, 86–110.

Mistrik, R., Aligizakis, N., Schymanski, E. and Williams, A. (2019) S19 | MZCLOUD | mzCloud Compounds (Version NORMAN-SLE-S19.0.2.0).

Mullard, G., Allwood, J.W., Weber, R., Brown, M., Begley, P., Hollywood, K.A., Jones, M., Unwin, R.D., Bishop, P.N., Cooper, G.J.S. and Dunn, W.B. (2015) A new strategy for MS/MS data acquisition applying multiple data dependent experiments on Orbitrap mass spectrometers in non-targeted metabolomic applications. Metabolomics 11, 1068–1080.

O’Hagan, S., Dunn, W.B., Brown, M., Knowles, J.D. and Kell, D.B. (2005) Closed-Loop, Multiobjective Optimization of Analytical Instrumentation: Gas Chromatography/Time-of-Flight Mass Spectrometry of the Metabolomes of Human Serum and of Yeast Fermentations. Analytical Chemistry 77, 290–303.

O’Hagan, S. and Kell, D.B. (2017) Consensus rank orderings of molecular fingerprints illustrate the ‘most genuine’ similarities between marketed drugs and small endogenous human metabolites, but highlight exogenous natural products as the most important ‘natural’ drug transporter substrates. ADMET & DMPK 5, 85–125.

O’Hagan, S., Wright Muelas, M., Day, P.J., Lundberg, E. and Kell, D.B. (2018) GeneGini: Assessment via the Gini Coefficient of Reference "Housekeeping" Genes and Diverse Human Transporter Expression Profiles. Cell Syst 6, 230–244 e1.

Psychogios, N., Hau, D.D., Peng, J., Guo, A.C., Mandal, R., Bouatra, S., Sinelnikov, I., Krishnamurthy, R., Eisner, R., Gautam, B., Young, N., Xia, J., Knox, C., Dong, E., Huang, P., Hollander, Z., Pedersen, T.L., Smith, S.R., Bamforth, F., Greiner, R., McManus, B., Newman, J.W., Goodfriend, T. and Wishart, D.S. (2011) The human serum metabolome. PLoS One 6, e16957.

Sansone, S.-A., Fan, T., Goodacre, R., Griffin, J.L., Hardy, N.W., Kaddurah-Daouk, R., Kristal, B.S., Lindon, J., Mendes, P., Morrison, N., Nikolau, B., Robertson, D., Sumner, L.W., Taylor, C., van der Werf, M., van Ommen, B., Fiehn, O. and Members, M.S.I.B. (2007) The Metabolomics Standards Initiative. Nature Biotechnology 25, 846–848.

Schymanski, E.L., Jeon, J., Gulde, R., Fenner, K., Ruff, M., Singer, H.P. and Hollender, J. (2014) Identifying Small Molecules via High Resolution Mass Spectrometry: Communicating Confidence. Environmental Science & Technology 48, 2097–2098.

Sumner, L.W., Amberg, A., Barrett, D., Beale, M.H., Beger, R., Daykin, C.A., Fan, T.W.M., Fiehn, O., Goodacre, R., Griffin, J.L., Hankemeier, T., Hardy, N., Harnly, J., Higashi, R., Kopka, J., Lane, A.N., Lindon, J.C., Marriott, P., Nicholls, A.W., Reily, M.D., Thaden, J.J. and Viant, M.R. (2007) Proposed minimum reporting standards for chemical analysis. Metabolomics 3, 211–221.

Superti-Furga, G., Lackner, D., Wiedmer, T., Ingles-Prieto, A., Barbosa, B., Girardi, E., Goldmann, U., Gurtl, B., Klavins, K., Klimek, C., Lindinger, S., Lineiro-Retes, E., Muller, A.C., Onstein, S., Redinger, G., Reil, D., Sedlyarov, V., Wolf, G., Crawford, M., Everley, R., Hepworth, D., Liu, S., Noell, S., Piotrowski, M., Stanton, R., Zhang, H., Corallino, S., Faedo, A., Insidioso, M., Maresca, G., Redaelli, L., Sassone, F., Scarabottolo, L., Stucchi, M., Tarroni, P., Tremolada, S., Batoulis, H., Becker, A., Bender, E., Chang, Y.N., Ehrmann, A., Muller-Fahrnow, A., Putter, V., Zindel, D., Hamilton, B., Lenter, M., Santacruz, D., Viollet, C., Whitehurst, C., Johnsson, K., Leippe, P., Baumgarten, B., Chang, L., Ibig, Y., Pfeifer, M., Reinhardt, J., Schonbett, J., Selzer, P., Seuwen, K., Bettembourg, C., Biton, B., Czech, J., de Foucauld, H., Didier, M., Licher, T., Mikol, V., Pommereau, A., Puech, F., Yaligara, V., Edwards, A., Bongers, B.J., Heitman, L.H., AP, I.J., Sijben, H.J., van Westen, G.J.P., Grixti, J., Kell, D.B., Mughal, F., Swainston, N., Wright-Muelas, M., Bohstedt, T., Burgess-Brown, N., Carpenter, L., Durr, K., Hansen, J., Scacioc, A., Banci, G., Colas, C., Digles, D., Ecker, G., Fuzi, B., Gamsjager, V., Grandits, M., Martini, R., Troger, F., Altermatt, P., Doucerain, C., Durrenberger, F., Manolova, V., Steck, A.L. et al. (2020) The RESOLUTE consortium: unlocking SLC transporters for drug discovery. Nat Rev Drug Discov.

Tautenhahn, R., Cho, K., Uritboonthai, W., Zhu, Z., Patti, G.J. and Siuzdak, G. (2012) An accelerated workflow for untargeted metabolomics using the METLIN database. Nature Biotechnology 30, 826–828.

Treutler, H., Tsugawa, H., Porzel, A., Gorzolka, K., Tissier, A., Neumann, S. and Balcke, G.U. (2016) Discovering Regulated Metabolite Families in Untargeted Metabolomics Studies. Analytical Chemistry 88, 8082–8090.

Vaidyanathan, S., Broadhurst, D.I., Kell, D.B. and Goodacre, R. (2003) Explanatory Optimization of Protein Mass Spectrometry via Genetic Search. Analytical Chemistry 75, 6679–6686.

Vinaixa, M., Schymanski, E.L., Neumann, S., Navarro, M., Salek, R.M. and Yanes, O. (2016) Mass spectral databases for LC/MS-and GC/MS-based metabolomics: State of the field and future prospects. TrAC Trends in Analytical Chemistry 78, 23–35.

Wright Muelas, M., Mughal, F., O’Hagan, S., Day, P.J. and Kell, D.B. (2019) The role and robustness of the Gini coefficient as an unbiased tool for the selection of Gini genes for normalising expression profiling data. Scientific Reports 9, 17960.

Zelena, E., Dunn, W.B., Broadhurst, D., Francis-McIntyre, S., Carroll, K.M., Begley, P., O’Hagan, S., Knowles, J.D., Halsall, A., Consortium, H., Wilson, I.D. and Kell, D.B. (2009) Development of a robust and repeatable UPLC-MS method for the long-term metabolomic study of human serum. Anal Chem 81, 1357–64.

