## Supplementary Information 1: Supplementary Methods, Figures and Tables for "An untargeted metabolomics strategy to measure differences in metabolite uptake and excretion by mammalian cell lines"

#### Incubation of cells in serum

Once cell counts were performed on harvested cells, an appropriate volume for the required number of cells was transferred to 2 mL Eppendorf tubes. Cells were pelleted by centrifugation at 2,000 rpm for 5 minutes, supernatant discarded, and cells washed with pre-warmed (37 ^o^C) DBPS without calcium or magnesium. Following cell pelleting by centrifugation (2,000 rpm for 5 minutes) cells were resuspended in 200 µL of pooled human serum (BioIVT, Lot BRH1413770, Cat: HMSRM, mixed gender 0.1 um filtered) and transferred to a shaking incubator set at 200 rpm for the required time.

At each time point, a 10 µL was taken from the cell suspension for cell count and diameter measurements. Cells were centrifuged at 2,000 rpm for 5 minutes, supernatant collected and immediately stored on dry ice. The remaining pellet was firstly washed by resuspending in 400 µL of prewarmed (37 ^o^C) DPBS. This cell suspension was then centrifuged (2,000 rpm for 5 minutes) and supernatant discarded. To the cell pellets then 400 µL of cold 80 % (v/v) methanol was added and pipetted vigorously to break open cells and release intracellular metabolites. The mixture was subjected to three cycles of snap freezing in liquid nitrogen followed by thawing on ice after which the samples were centrifuged at 13,300 rpm for 10 mins at 4 °C for 15 minutes. The supernatant was collected and stored on dry ice. The remaining cell pellet was then gently resuspended in 200 µL TRIzol (Invitrogen, Cat no: 11596026) and stored immediately on dry ice. All samples were subsequently stored at -80 °C until preparation for analysis.

#### MS acquisition settings

Full-scan MS data was acquired in the Orbitrap mass analyser in the m/z range 66.7-1,000 with a mass resolution of 70,000 Full Width Half Maximum (FWHM) at m/z = 200, a chromatographic peak width to 4 s (FWHM), AGC target = 1 x 10^6^ and maximum injection time = 100 ms. HESI source and ion transfer parameters applied were as follows: Sheath flow rate (arbitrary units) = 50, auxiliary gas flow rate (arbitrary units) = 12, sweep gas flow rate (arbitrary units) = 2, spray voltage = 3.5 kV (positive) and -2.8 kV (negative), capillary temperature (°C) = 263, S-lens RF level (%) = 58.5, auxiliary gas heater temperature (°C) 425, source position = C.

Data-dependent MS/MS (ddMS^2^) data acquisition was performed on pooled samples at the end of the analytical run as described in guidelines by (Broadhurst *et al.*, 2018). In a manner similar to that proposed by (Mullard *et al.*, 2015), ddMS^2^ data acquisition was performed on 4 precursor mass ranges in triplicate injections: 66.7-1,000, 66.7-300, 300-600 and 600-900. Data was acquired in the orbitrap mass analyser with a mass resolution of 17,500 Full Width Half Maximum (FWHM) at m/z = 200 and a chromatographic peak width to 4 s (FWHM), AGC target = 1 x 10^5^, maximum injection time = 54 ms, loop count = 5, isolation window = 1.2 m/z, stepped (N)CE = 20, 50 and 80, minimum AGC target 1 x 10^3^, exclude isotopes = on and dynamic exclusion = 6.0 s.

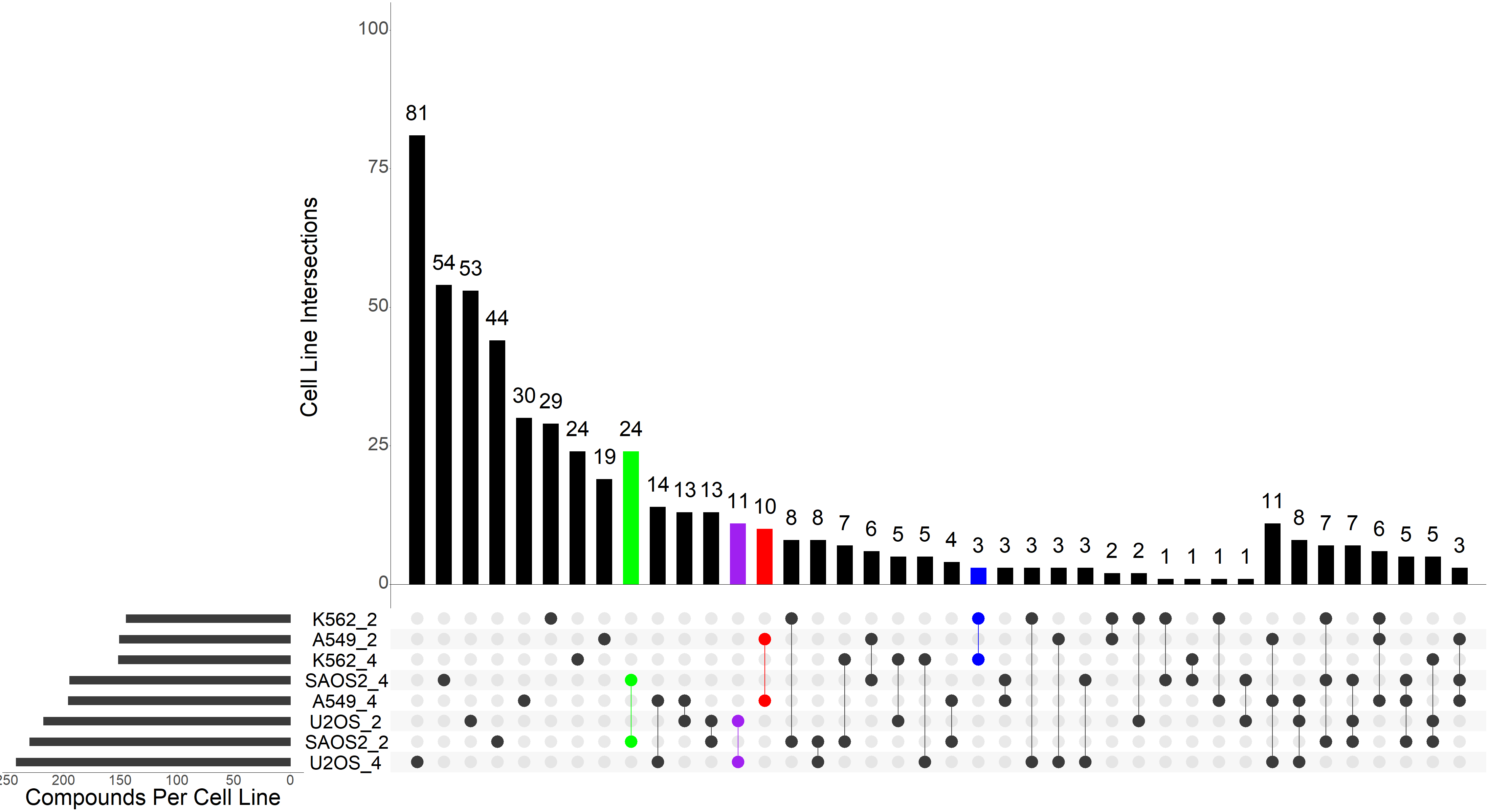

Shared and unique excreted serum compounds

Shared and unique consumed serum compounds

A

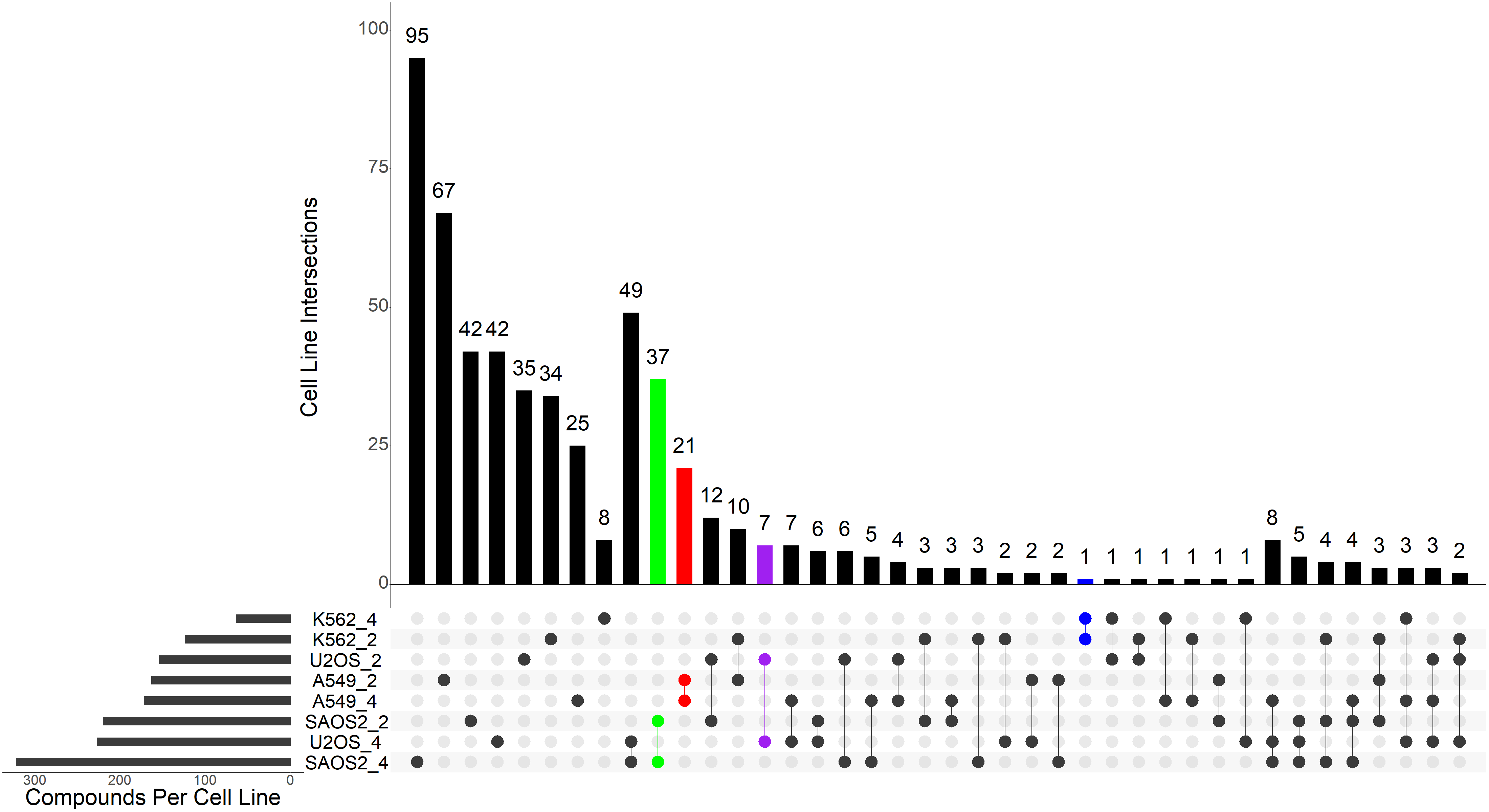

B

Supplementary Figure 1. UpSet plot of shared and unique A) consumed or B) excreted metabolites between cell lines and densities using compounds with Log_2_ FC < -0.5 or > 0.5 and P < 0.05 (t-test) . Cell line unique consumed compounds coded in colour: A549, red; K562, blue; SAOS2, green; U2OS, purple.

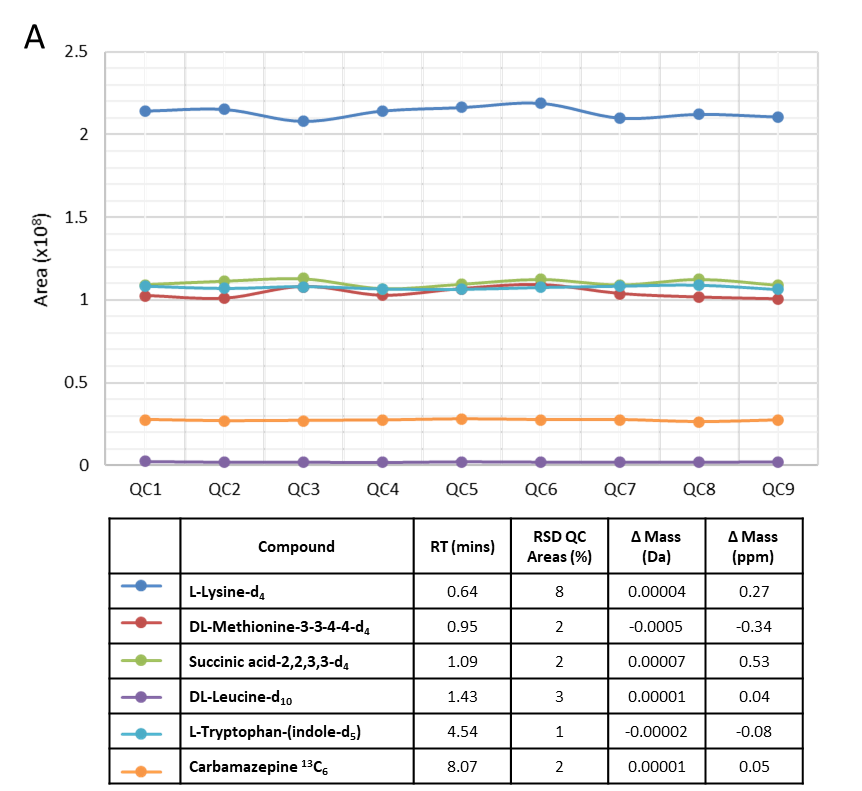

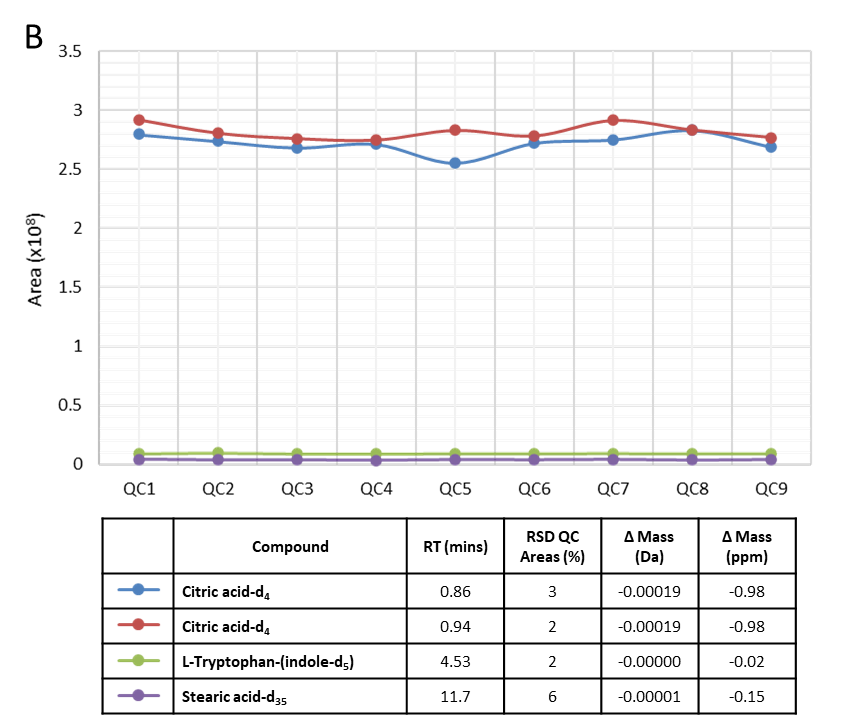

Supplementary Figure 2 Peak areas, retention time (RT), % RSD and mass accuracy of detected spiked stable isotope labelled standards in QC samples. A) ESI+ and B) ESI-.

Supplementary Table 1: Summary of ESI+ LC-MS/MS results obtained following preprocessing using CD3.1 for spent serum analysis following incubation with 4 different cell lines.

|  |  |  | **Number** | **Percentage** | |
| --- | --- | --- | --- | --- | --- |
|  |  | **Features** | 244,132 |  |  |
|  |  | **Compounds** | 8,663 |  |  |
|  |  | **Background compounds** | 882 | 10.2 | % of total compounds |
|  |  | **Other excluded compounds** | 2,957 | 34.1 |  |
| **Sample compounds** |  | **Total** | 4,824 | 55.7 |  |
|  |  | **Total with % QC CV < 15** | 3,822 | 79.2 | % total sample compounds |
|  |  | **No MS2 spectra** | 2,582 | 53.5 |  |
|  |  | **MS2** | 2,242 | 46.5 |  |
|  |  | **MS2 for prefered ion** | 2,072 | 92.4 | % of MS2 |
|  |  | **MS2 for other ion** | 170 | 7.6 |  |
|  | **Annotation and Identification** | **Full match to Proposed Molecular Formula (level 4)** | 3,959 | 82.1 | % total sample compounds |
|  |  | **No match to Proposed Molecular Formula** | 627 | 13.0 |  |
|  |  | **Full match to ChemSpider** | 1,672 | 34.7 | % total sample compounds |
|  |  | **Full match Proposed Molecular Formula and ChemSpider (level 3)** | 1,503 | 31.2 | % total sample compounds |
|  |  | **Full match to Mass List** | 782 | 16.2 | % total sample compounds |
|  |  | **Full match Proposed Molecular Formula and Mass Lists (level 3)** | 664 | 84.9 | % Mass List Matches |
|  |  | **mzCloud Best Match ≥70% score** | 398 | 8.3 | % total sample compounds |
|  |  | **Full match Proposed Molecular and mzCloud Best Match ≥70% score (level 2)** | 158 | 39.7 | % mzCloud Match ≥70% score |

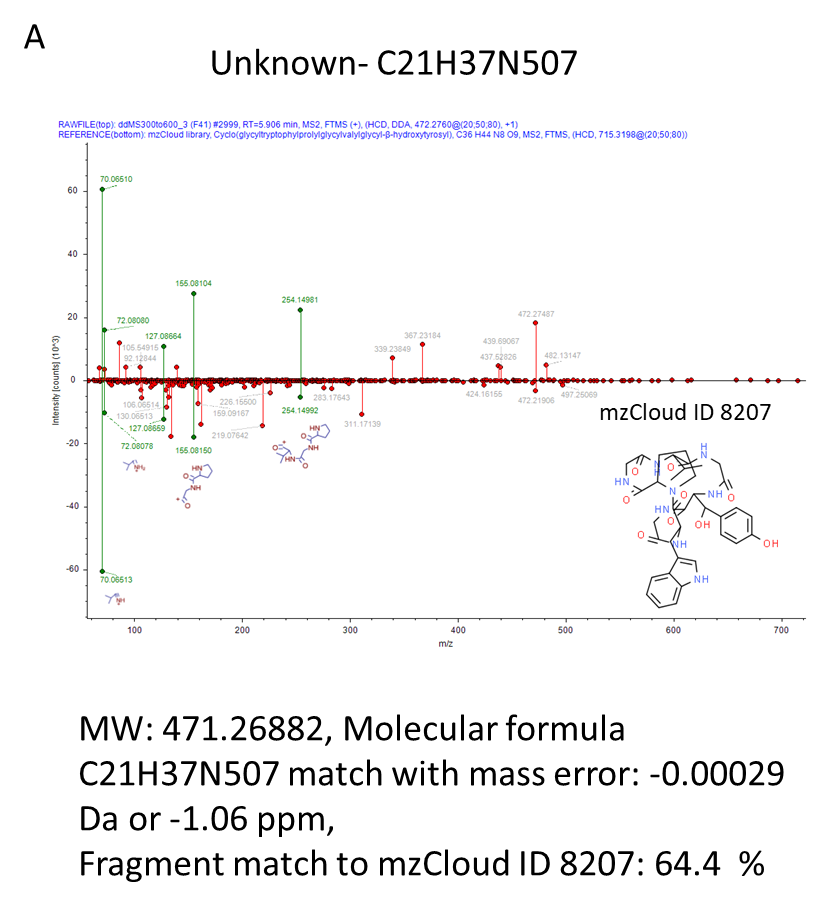

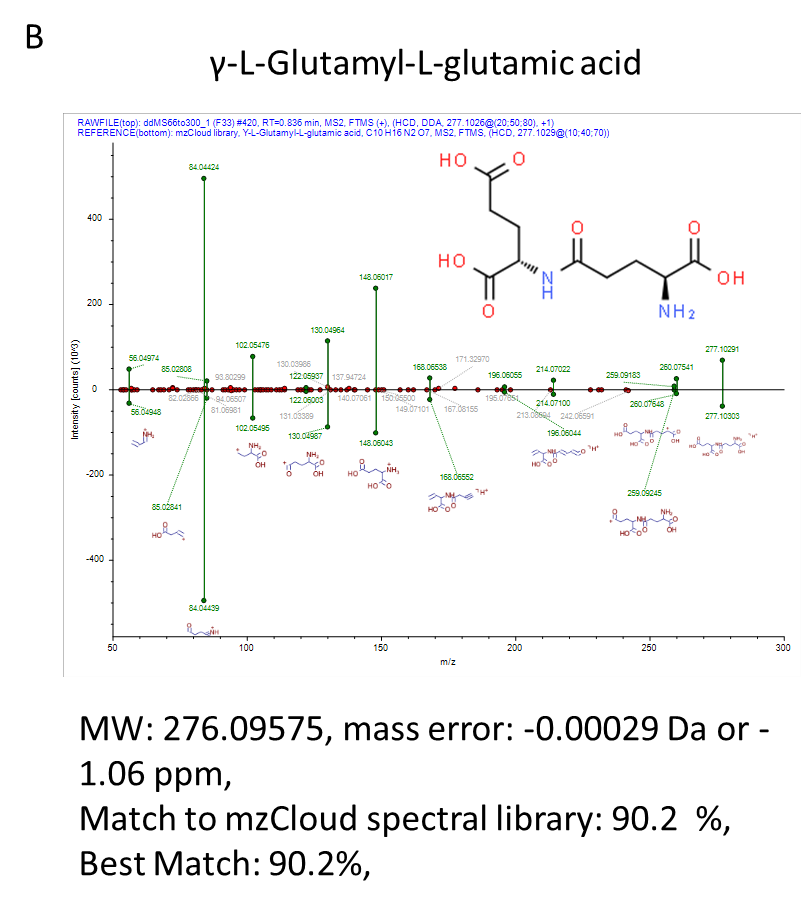

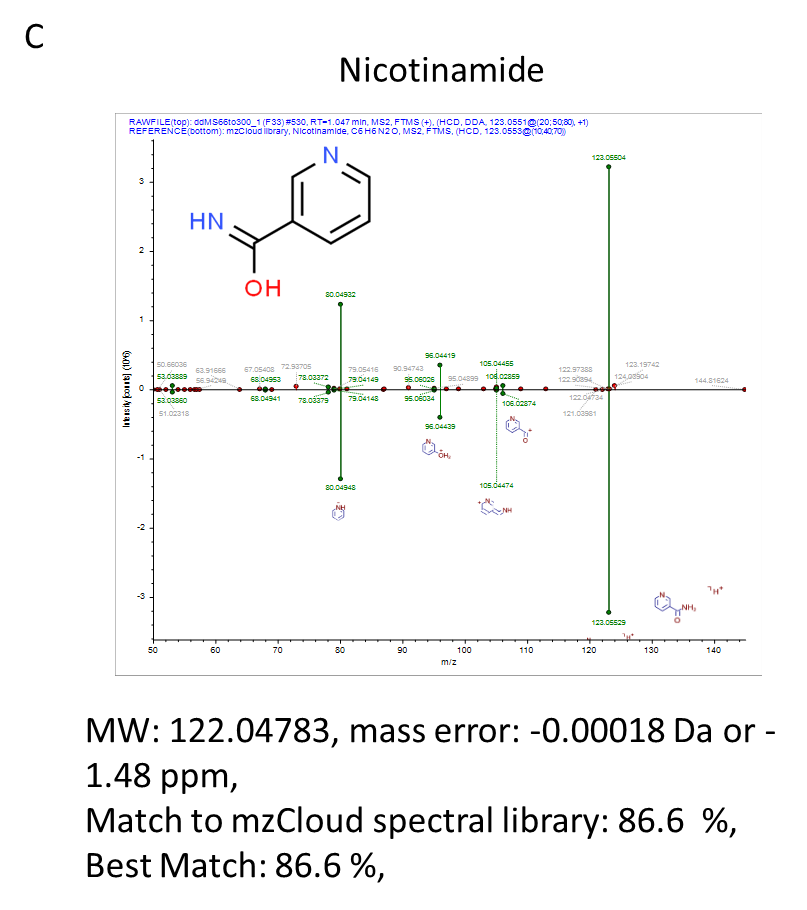

Supplementary Figure 3: Spectral matching of selected consumed and/or secreted compounds against mzCloud A) Unknown with match to molecular formula C21H37N507, B) γ-L-Glutamyl-L-glutamic acid and C) Nicotinamide

### References

Broadhurst, D., Goodacre, R., Reinke, S.N., Kuligowski, J., Wilson, I.D., Lewis, M.R. and Dunn, W.B. (2018) Guidelines and considerations for the use of system suitability and quality control samples in mass spectrometry assays applied in untargeted clinical metabolomic studies. *Metabolomics* **14,** 72.

Mullard, G., Allwood, J.W., Weber, R., Brown, M., Begley, P., Hollywood, K.A., Jones, M., Unwin, R.D., Bishop, P.N., Cooper, G.J.S. and Dunn, W.B. (2015) A new strategy for MS/MS data acquisition applying multiple data dependent experiments on Orbitrap mass spectrometers in non-targeted metabolomic applications. *Metabolomics* **11,** 1068-1080.
